# Using green space virtual reality to prevent stress-induced working memory impairment

**DOI:** 10.1101/2024.12.31.630298

**Authors:** Yingying Xu, Zixuan Yang, Xiaoxuan Zhou, Qingyi Yao, Guohang Dai, Wei Liu, Hui Zhao

**Author notes:** **Correspondence:** Dr. Hui Zhao State Key Laboratory of Cognitive Neuroscience and Learning, Beijing Normal University Address: No. 19, Xinjiekouwai Street, Beijing, China, 100875 Dr. Wei Liu School of Psychology, Central China Normal University Address: the 8^th^ floor, Nanhu Complex Building, No 152 Luoyu Road, Wuhan. Postal Code: 430079.

## Abstract

Exposure to green space is associated with both physical and mental health benefits, including the potential to buffer acute stress responses, positioning it as a promising non-pharmacological approach to protect cognitive functions against stress. However, urban residents often face significant barriers in accessing green spaces, which are not equitably distributed. Virtual Reality (VR) technology offers a potential solution by simulating green space that could be made accessible to a broader demographic. Here, we explored whether VR-stimulated green space could dampen biological stress responses (i.e., heart rate and cortical response) and prevent stress-induced working memory (WM) impairment. Healthy young participants underwent acute stress induction followed by 15 minutes of VR-based green space (N=36) or control empty space (N=30) intervention. Although we observed the expected stress-induced increase in heart rates and elevated cortisol levels under stress, VR green space exposure failed to temper cortisol responses compared to the control VR space. Further, VR green space exposure did not bring benefits to protect working memory performance under stress across three WM tasks. Applying a Bayesian analysis approach throughout enabled us to find substantial evidence for the absence of an effect of VR-based green space exposure on biological markers of acute stress responses and working memory performance. Our findings suggest that VR-generated green space may not effectively replicate the stress-buffering effects of actual green space exposure. We discussed the implications of our findings regarding the potential and limitations of using VR or green space exposure to buffer stress responses.

## Introduction

Exposure to acute stress influences a broad spectrum of cognitive functions in humans^1^, such as attention^2,3^, cognitive control^4,5^, social cognition^6^, and decision making^7^. Among these, working memory (WM) is notably vulnerable to acute stress and related stress hormones^8–10^. This vulnerability is mirrored by similar deficits seen in stress-related psychiatric disorders, such as depression^11^ and PTSD^12^. The activation of the hypothalamic-pituitary-adrenal (HPA) axis by acute stress, which orchestrates the release of glucocorticoids, is believed to be the underlying mechanism for these stress-induced working memory impairments^10,13,14^. Neuroscientific evidence in humans have shown that acute stress decreases frontal theta oscillation^15^ and reduces activity in the dorsolateral prefrontal cortex (DLPFC) during the working memory task^16^. Understanding the molecular and neural mechanisms of stress-induced working memory impairments is crucial for developing interventions to protect working memory in healthy individuals facing stressful situations and to aid recovery in individuals with stress-related psychiatric disorders. In the present study, we aim to prevent stress-induced working memory impairments directly by reducing biological stress responses. Using Virtual Reality (VR) technology and insights from environment psychology and neuroscience^17^, we developed the VR-simulated green space intervention aimed at reducing stress responses and protect working memory against stress.

A growing body of research has demonstrated psychological benefits of exposure to natural environment, including but not limited to improved cognitive functions^18,19^, reduced negative emotions and stress^20,21^. Empirical evidence from a study involving approximately 5,000 UK residents revealed buffered declined mental health during the COVID-19 pandemic^22^. Critically, nature’s beneficial effects on reducing stress are evident not only through subjective or cognitive evaluations, but also via physiological and neural markers of stress, such as reductions in heart rate, blood pressure, cortisol levels (a stress-related hormone), and amygdala activity under stress conditions ^20,23^. As an alternative of natural environment, however, the availability of green spaces remains constrained for many, especially urban dwellers. A recent systematic review highlighted an increase in the use of private gardens across the examined studies, while 77% of the studies reported a decrease in the use of public parks^24^, highlighting the challenge of equitable green space access. The advent of virtual reality (VR) offers a promising avenue by simulating green environments that partly replicated stress-reducing effects observed in natural settings^25–32^. However, despite accumulating evidence on its efficacy in reducing negative emotions, physiological responses to VR immersion in nature show variability^33^, suggesting that the application of VR immersion in nature for therapeutic purposes requires further exploration and validation.

Moreover, compared to reducing biological stress responses, protecting cognitive functions under stress remains a critical, yet less explored, objective. Various strategies have been undertaken to counteract stress-induced cognitive impairments^34^. Emotional strategies, such as recalling positive memories^35^ and goal-shifting^36^ have been shown to reduce cortisol responses to stress. However, their effectiveness in protect cognitive functions under stress is untested or not evident, likely due to the cognitive resources required for these emotional strategies. Memory retrieval impairments under stress can be prevented either by pharmacologically inhibiting the biological responses to acute stress^37,38^ or by enhancing memory representations through post-encoding retrieval practice^39^. Non-invasive brain stimulation targeting the DLPFC has been shown to effectively prevent stress-induced working memory deficits in humans^40^. While these pharmacological and brain stimulation interventions may prove effective, their applicability in real-life settings, such as classrooms, and among broader populations, including children and older adults, presents significant challenges. Consequently, it is crucial to explore alternative, safe methods for preventing stress-related working memory deficits in humans. Research has demonstrated that exposure to virtual green spaces yields restorative effects, including improved emotional well-being and reduced biological stress responses. To our knowledge, no study has yet used VR-stimulated green space to buffer the negative effects of stress on cognitive functions. In this study, given the need for non-pharmacological and non-invasive solutions to protect working memory under stress, and considering the potential of green space VR, we specifically explored whether VR green space could prevent stress-induced working memory impairments.

The present study investigated the potential of using VR-simulated green space to diminish biological responses (i.e., heart rate and cortisol rise) and cognitive consequences (specifically, impairments in work memory) of acute stress (**Figure1**). We exposed healthy participants to an acute stressor before the VR session, which involved exposure to either a simulated green space (N=36) or size-matched empty control space (N=30). Salivary cortisol samples were collected at three time points: baseline (SC0), immediately post-VR immersion (SC1=SC0+50 minutes), and at the study’s end (SC2=SC0+80 minutes). Similarly, heart rates were recorded at three time points: baseline (HR0), post-stress induction (HR1=HR0+30 minutes), and immediately post-VR immersion (HR2=HR0+50 minutes). By analyzing the temporal dynamics of biological stress markers, we confirmed that our stress induction protocol successfully triggered the rapid release of corticosteroids and activated the HPA axis, as evidenced by increased heart rate and cortisol levels. These two physiological markers were subsequently analyzed across the two group conditions to explore the effect of VR green space on stress. Additionally, working memory performance was accessed pre-stress induction and post-VR immersion using three different paradigms (i.e., Digit Symbol Substitution Task (DSST), visuo-spatial (VS) 3-back task, and the phonological loop (PL) 3-back task). We hypothesized that exposure to VR-simulated green space, compared to the VR-immersed control empty space, would buffer the effects of acute stress by (1) dampen biological stress responses and (2) protect working memory against stress.

**Figure 1.**
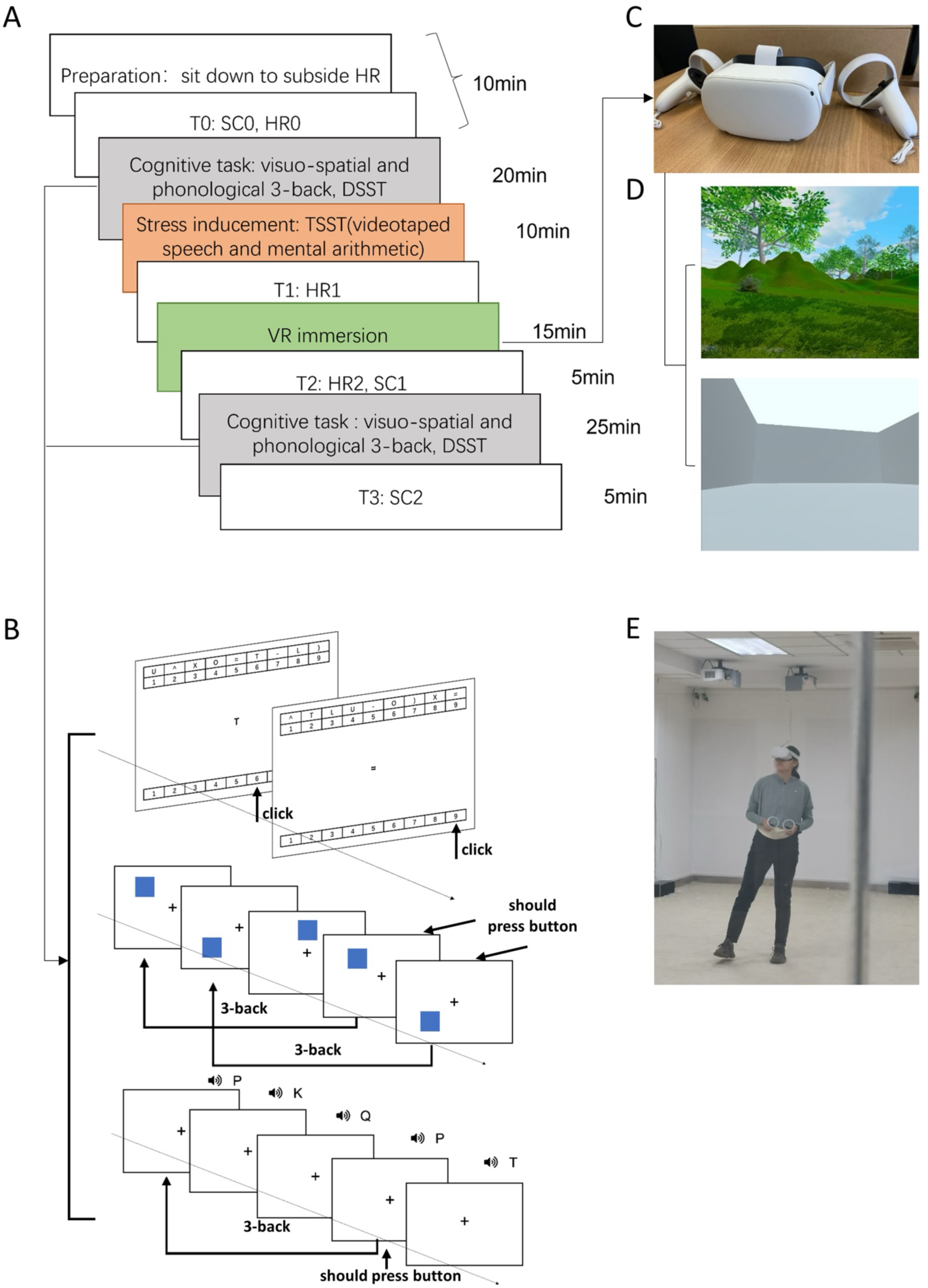
Schematic of the Experimental Procedure. (A) All participants underwent the identical experimental protocol, differing only in the VR immersion scene—either a natural green space or a gray control space. We assessed biological responses to acute stress using neuroendocrine and physiological measures, specifically salivary cortisol (SC) and heart rate (HR). SC levels were measured at three time points: baseline (SC0), post-intervention (SC1), and post-task (SC2). Similarly, HR was recorded at baseline (HR0), before intervention (HR1), and post-intervention (HR2). Three distinct working memory tasks were administered: a visuo-spatial 3-back task (VS), a phonological loop 3-back task (PL), and the Digit Symbol Substitution Test (DSST). These assessments occurred immediately before stress induction and five minutes following the VR intervention. Acute stress was induced using the Trier Social Stress Test (TSST). (B) Working Memory Task: The tasks presented from top to bottom are the DSST, VS, and Phonological Loop (PL) tasks, respectively (see Section 2.5 for more details). (C) Virtual Reality Headset: Participants wore a virtual reality headset and used two handheld controllers to interact with the virtual environment (see Section 2.6 for more details). (D) Participant Views During VR Intervention: Visual perspectives of participants in the two groups during the VR intervention. The upper panel shows the green nature group, and the lower panel shows the control group. (E) Participant Interaction with the VR Scene. Images showing a participant exploring the VR-simulated environment. *Note: This photograph was provided by one of the authors for demonstration purposes and does not depict a real participant. Informed consent has been obtained from the individual for the use of this image in the paper. While this photo was not taken during the actual data collection, it was captured in the same environment using identical VR equipment*.

## 2. Methods

### 2.1 Participants

Sixty-six healthy, right-handed university students (aged 18-24; 55 females) with normal or corrected-to-normal vision participated in this study. All participants reported no history of neurological, psychiatric, or endocrine disorders, nor any current use of psychoactive drugs or corticosteroids. We obtained written informed consent prior to the experiment, and the University’s Institutional Review Board for Human Subjects approved the study protocols. We collected demographic data, including age, gender, and major, to control for the effects of the participants’ prior knowledge on the experimental procedure. The participants were randomly assigned to either the Natural Green Scene VR group (n_nature_=36, aged 20.07±1.46 years, 32 females) or the Empty Gray Control group (n_control_=30, aged 19.87±1.23 years, 23 females). We observed no significant differences in age (t=0.09, *p*=0.92) or gender distribution (χ²=0.99, *p*=0.32) between the groups.

### 2.2 General Procedure

All experimental procedures were conducted between 2:00 p.m. and 6:00 p.m. to control for diurnal variations in cortisol secretion. Upon arrival, participants rested for 5-10 minutes to ensure accurate baseline heart rate measurements. Participants were instructed to avoid fluid intake for 30 minutes before the study began, facilitating accurate acute cortisol level assessments. This study investigated the effectiveness of VR-simulated green spaces in mitigating the biological and cognitive effects of acute stress. We enrolled a healthy cohort, randomly assigning participants to one of two conditions: exposure to a VR-simulated green space or a comparable empty control scene. Initially, participants underwent baseline assessments (T0) of heart rate and salivary cortisol (see *Physiological Measurements of Stress* below) along with working memory performance (see *Working Memory Tasks* below). Subsequently, they underwent a stress induction procedure (see *Stress Induction* below). Immediately following stress induction (T1), heart rate measurements were taken as an immediate response to the stressor. During the stress recovery phase, participants were exposed to their assigned VR environments (Nature or Control). Post-VR session (T2), the second heart rate and salivary cortisol measurements were taken. Participants then repeated all working memory tasks, and finally, a third salivary sample (T3) was collected as an indicator of slow recovery from stress.

### 2.3 Stress induction

We employed a modified version of the Trier Social Stress Test (TSST)^41^ to induce acute stress in a laboratory setting. Previous research has demonstrated that this paradigm consistently provokes an increase in both hypothalamic-pituitary-adrenal (HPA) axis and sympathetic nervous system (SNS) activity^42^. Our adaptation includes a videotaped free speech, and a mental quick arithmetic task performed in front of a panel of experimenters who maintain a reserved demeanor, lasting a total of 10 minutes. In a deviation from the original, we replaced subtraction problems with two-digit multiplication challenges (e.g., 56×47), issuing an alarm for incorrect or delayed responses within a 5-second window. Immediately after that, participants were asked to read a textbook section and then simulate a teaching role, akin to a high school teaching certification interview. This modification is designed to intensify mental stress compared to the original subtraction tasks. Prior to its implementation, we confirmed that completing these calculations within the allotted time was nearly unfeasible.

### 2.4 Physiological measurements of stress

We employed heart rate (HR) to evaluate the autonomic nervous system’s response and measured cortisol levels to assess the hypothalamic-pituitary-adrenal (HPA) axis’s reaction to stress. HR data were collected using a Mi NFC 7 sports wristband and recorded continuously throughout the experimental protocol. We calculated the mean HR for specific time windows as defined by different time points illustrated in **Figure 1**.

Cortisol measurements were taken at baseline, immediately after VR exposure, and following the second cognitive task. Salivary samples (2 mL) were collected using cryogenic tubes (Sarstedt, Suzhou, China). Participants were instructed to hold saliva in their mouths for 3 minutes before collection and to refrain from eating, drinking, or smoking for 30 minutes prior to arrival. All samples were stored at -80°C for a maximum of 3 months before analysis. Salivary-free cortisol concentrations were measured using an enzyme-linked immunosorbent assay (ELISA) with a kit (model 202401, Chundubio, Wuhan, China) by trained professionals.

### 2.5 Working Memory Task

We used the Digit Symbol Substitution Test (DSST) to evaluate attention components in working memory^43,44^. In a block design, participants completed nine trials per block. The correspondence rule between icons and numbers remained constant within each block and was carried over to the next block. In each trial, one of the nine icons was randomly displayed at the center of the screen without repetition (see Figure 1B, first panel). Participants were instructed to click the number corresponding to the given icon as quickly as possible using a mouse to proceed to the next trial. A practice session of nine trials was conducted before the formal task began. The DSST task lasted 90 seconds for the first cognitive task and 270 seconds for the second.

To evaluate working memory across different modalities, we conducted a 3-back task with visual (VS task) and auditory (PL task) stimuli^45,46^. Participants completed two or three cycles of 30 trials each (two cycles in the first cognitive task, three in the second), consisting of 10 target trials. Each trial began with a 1000 ms resting fixation, followed by a random stimulus lasting 500 ms. In the VS task, a blue square appeared randomly in one of eight locations around the resting fixation (three at the top and bottom, one at the left and right; see Figure 1B, second panel). Participants were instructed to indicate whether the current location matched the location three trials back by pressing a button with their index finger. An 18-trial practice session with six target trials was provided before the main task. The PL task included an 18-trial practice session (with six target trials) and followed the same cycle structure as the VS task. The stimuli were random letters that were easily distinguishable; all auditory stimuli were normalized and presented for 500 ms. Other parameters and instructions were the same as in the VS task.

### 2.6 Virtual reality space exposure

To simulate virtual reality (VR) green space and blank control exposure in the current experiment, we generated two size-matched VR environments without sound. Each scene covered an area of 100 × 100 square meters, with consistent light source properties. For the green space group, a grassy valley surrounded by five hills was created using Unity and Unreal Engine 4 (UE4). The hills were dotted with ten trees and various colorful flowers. The control group’s scene consisted solely of four enclosed grey walls and a grey floor (see Fig. 1D). All VR environments were static to avoid introducing any stressors. Participants viewed the VR scenes through an Oculus Quest 2 headset (Meta, Inc., Menlo Park, CA, USA) and could explore the environment freely using handheld controllers. They could interact with the scene by touching features, climbing hills, and walking within a 10 × 15 square meter area (see Fig. 1E). Prior to VR exposure, participants received brief instructions on using the controllers. All participants started at the center of the scene.

### 2.7 Statistical analyses

In this study, we employed both traditional Null Hypothesis Significance Testing (NHST) and Bayesian data analysis^47^ to evaluate the effects of VR-simulated green spaces on stress responses and working memory performance. We analyzed five dependent variables independently: Heart Rate (HR), Cortisol Secretion (SC), Reaction Time (RT) of DSST, Accuracy (ACC) of the PL and ACC of the VS. During NHST, we assessed stress induction through repeated measure ANOVA (RMANOVA) at various testing points and explored group differences using two-factor ANOVA, with group as the between-group variable. Comparisons of the area under the curve (AUC) for HR and SC between groups were conducted using a two-sample t-test. For the Bayesian analyses, we used Bayes Factors (BF)^48^ to assess the data. A BF_10_ greater than 3 indicates support for the alternative hypothesis, a BF_10_ less than 1/3 supports the null hypothesis, and a BF_10_ between 1/3 and 3 indicates inconclusive evidence. In addition, we conducted two exploratory analyses using subsamples drawn from the entire participant pool. Each analysis involved repeated applications of NHST and Bayesian analysis to assess group differences in biological stress responses and WM performance.

### 2.8 Data and code availability

Behavioral and biological data, specifically HR and SC, are available on the Open Science Framework (OSF) (https://osf.io/3s46m/?view_only=6eaef2e33e474066a1971861432310eb). The source files for the VR scene employed in this study can be accessed in the same repository. Data analyses were conducted using SPSS (https://www.ibm.com/spss), R-4.3.2 (https://www.r-project.org/), and the JASP (https://jasp-stats.org/). Custom scripts for these analyses are also hosted on the same OSF repository.

## 3. Results

To examine the effects of VR stimulated green space exposure on reducing acute stress responses and preventing stress-induced impairments in working memory, we conducted four types of analyses: (1) manipulation checks to validate the efficacy of stress induction; (2) the influence of VR stimulated green space on the dynamics of heart rate and cortisol level following stress induction; (3) the effect of VR stimulated green space on working memory performance under stress; (4) exploratory analyses to examine individual differences in biological and cognitive outcomes of stress induction.

### 3.1 Manipulation checks of stress induction

To verify our stress induction, we analyzed the dynamic changes in heart rate and cortisol levels following the acute stress induction (**Figure 2**). We found significant time effect in both two measures (HR: F(2,128) = 26.03, p<.001, partial η^2^=0.29; SC: F(2,130) = 69.15, p<.001, partial η^2^=0.52). As an immediate response to acute stress, heart rate increased significantly from T0 to T1 (t= 5.41, *p* < .001, Cohen’s d =0.84; BF_10_ > 100), returning to baseline level at T3 (t= 1.43, *p* = .15, Cohen’s d =0.22; BF_10_= 0.41). As a slower response to acute stress, cortisol levels significantly increased from T0 to T2 (t = 11.75, *p* < .001, Cohen’s d = 1.90; BF_10_ > 100), and declined from T2 to T3 (t= 5.43, *p*<.001, Cohen’s d =0.88; BF_10_ > 100), but not subsided to resting level (t= 6.32, *p*<.001, Cohen’s d =1.02. BF_10_> 100). These dynamic patterns align with previous literature on stress induction in healthy humans, indicating successful stress induction^40,49,50^. Subsequently, we investigated the association between heart rate (HR) and cortisol secretion (SC) across participants. The Pearson correlation analysis revealed no significant relationship between the increase in HR and SC (r(65) = 0.09, *p* = .46, BF_10_ = 0.20), nor in the recovery phase (r(65) = 0.01, *p* = .93, BF_10_ = 0.16). These results suggest distinct response patterns between the neuroendocrine system and the autonomic nervous system.

**Figure 2.**
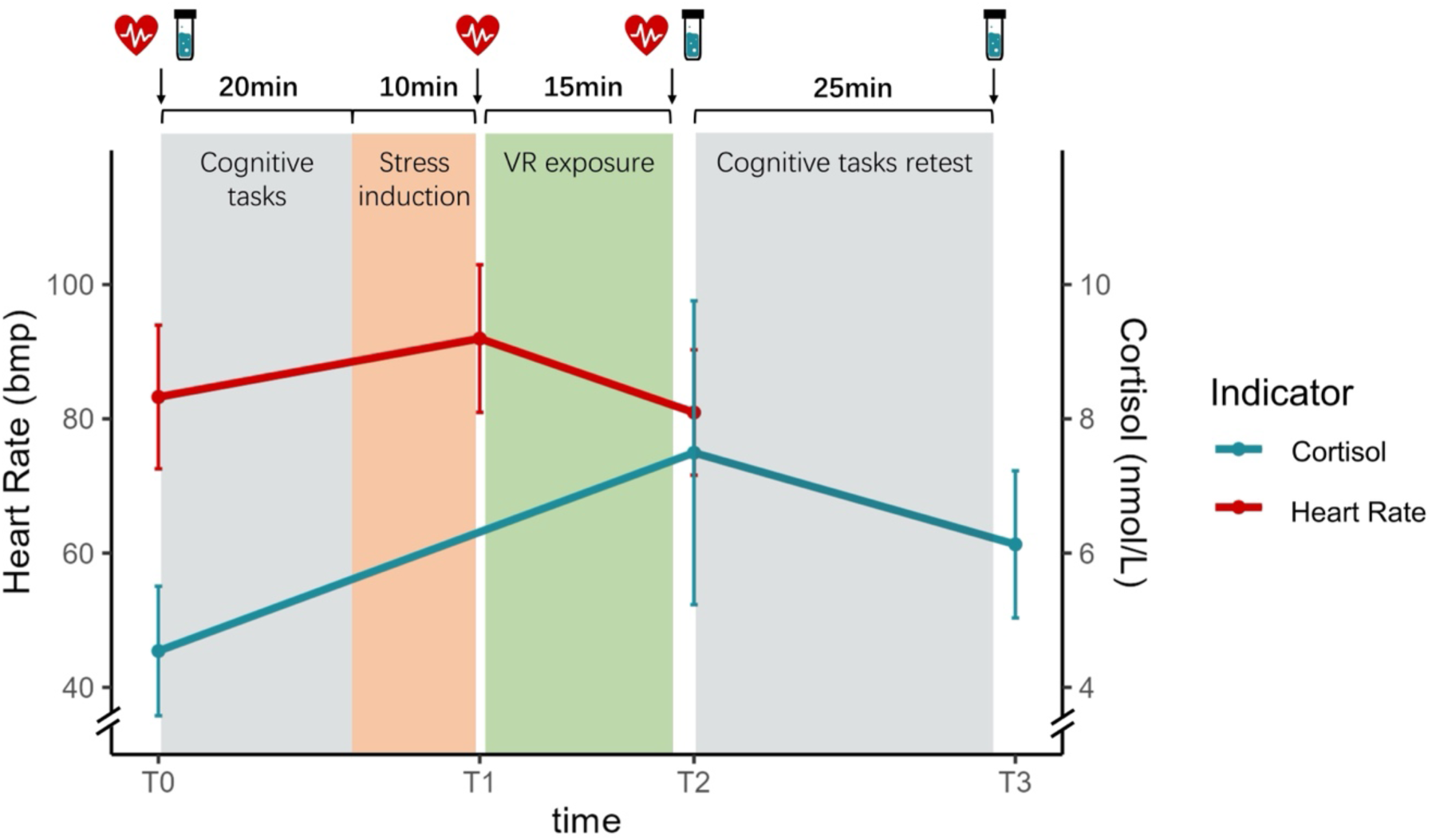
Dynamic changes in heart rate and cortisol levels confirm successful stress induction. In all participants, we observed a rapid and significant increase in heart rate (red line) from T0 (baseline) to T1 (pre-intervention), which returned to normal levels at T2 (post-intervention). Additionally, cortisol levels showed a gradual increase from T0 to T2, followed by a decrease from T2 to T3 (post-task). The gray shading indicates a 20–25-minute period during which participants engaged in cognitive tasks (i.e., working memory), the orange shading represents 10 minutes of acute stress induction, and the green shading denotes 15 minutes of VR intervention (natural green space or gray control scene).

### 3.2 Does VR stimulated green space dampen the biological stress responses?

We analyzed the between-group differences (Green vs. Control) in biological stress responses, specifically measuring heart rate and cortisol levels, to evaluate the dampening effect of VR-stimulated green spaces. Descriptive data for heart rate and cortisol at each time point are detailed separately for both groups in **Table 1**.

**Table 1.** Descriptive Statistics of Heart Rate and Cortisol Levels Across Time Points for Green and Control Groups.

| Indicator<sup>3</sup> | Timepoint/interval | Green(N=36) | Control(N=30) | NHST<sup>1</sup> |  |  | Bayesian<sup>2</sup> |
| --- | --- | --- | --- | --- | --- | --- | --- |
| | | Descriptive: Mean±SD | | F/t | p | $\eta_p^2/d$ | BF<sub>10</sub> |
| HR | T0 | 83.92±11.13 | 82.41±10.27 | 0.39 | .54 | 0.006 | 0.22 |
|  | T1 | 92.31±11.86 | 91.52±10.01 |  |  |  |  |
|  | T2 | 82.41±10.27 | 80.34±9.50 |  |  |  |  |
|  | T3 | NA<sup>4</sup> | NA |  |  |  |  |
| AUC<sub>HR</sub> | T0-T2 | 86.74±228.41 | 95.43±236.82 | 0.15 | .52 | 0.16 | 0.26 |
| SC | T0 | 4.61±1.00 | 4.46±0.93 | 0.004 | .95 | 0 | 0.18 |
|  | T1 | NA | NA |  |  |  |  |
|  | T2 | 7.41±2.43 | 7.60±2.08 |  |  |  |  |
|  | T3 | 6.13±1.05 | 6.13±1.16 |  |  |  |  |
| AUC<sub>SC</sub> | T0-T3 | 106.68±74.71 | 119.31±82.53 | 0.65 | .52 | 0.16 | 0.30 |

1. Null hypothesis significance testing (repeat measure ANOVA and independent t-test) were conducted to examine stress inducement (F, p values and partial $\eta^2$ ) and potential between-group difference in AUC (t, p values and Cohen' d) based on general data.
2. Bayesian analyses were performed to provide evidence for or against the hypothesis that virtual reality (VR) simulated scenes caused between-group differences in HR or SC (using Bayesian repeated measures ANOVA) or AUC (using Bayesian independent t-tests).
3. HR refers to heart rate, representing an immediate response to the stressor. SC stands for salivary cortisol, indicating a delayed response to the stressor. AUC, or area under the curve, estimates HR or SC changes over time, calculated using the trapezoidal method. The formula differs from individual curve models (see Supplementary Table 1 in the Supplementary Materials).
4. NA indicates that data was not collected at that time point.

Heart rate data analyzed using repeated measures ANOVA showed a significant main effect of time (F (2,126) = 25.45, *p*<.001, partial η^2^= 0.29), but no group differences (F(1,63)= 0.39, *p*= .54, partial η^2^= 0.006.) or time-group interactions (F(2,126) = 0.02, *p*= .88, partial η ^2^= 0.001; **Figure3A**). Bayesian repeated measures ANOVA confirmed the model including only time (BF_10_>100). The area under the curve (AUC) for heart rate from T0 to T2, calculated using the trapezoidal rule, showed no significant differences between groups (t(63)=0.15, p=.52, Cohen’s d=0.16; **Figure3C**), with Bayesian analysis providing strong support (BF_10_= 0.26).

**Figure 3.**
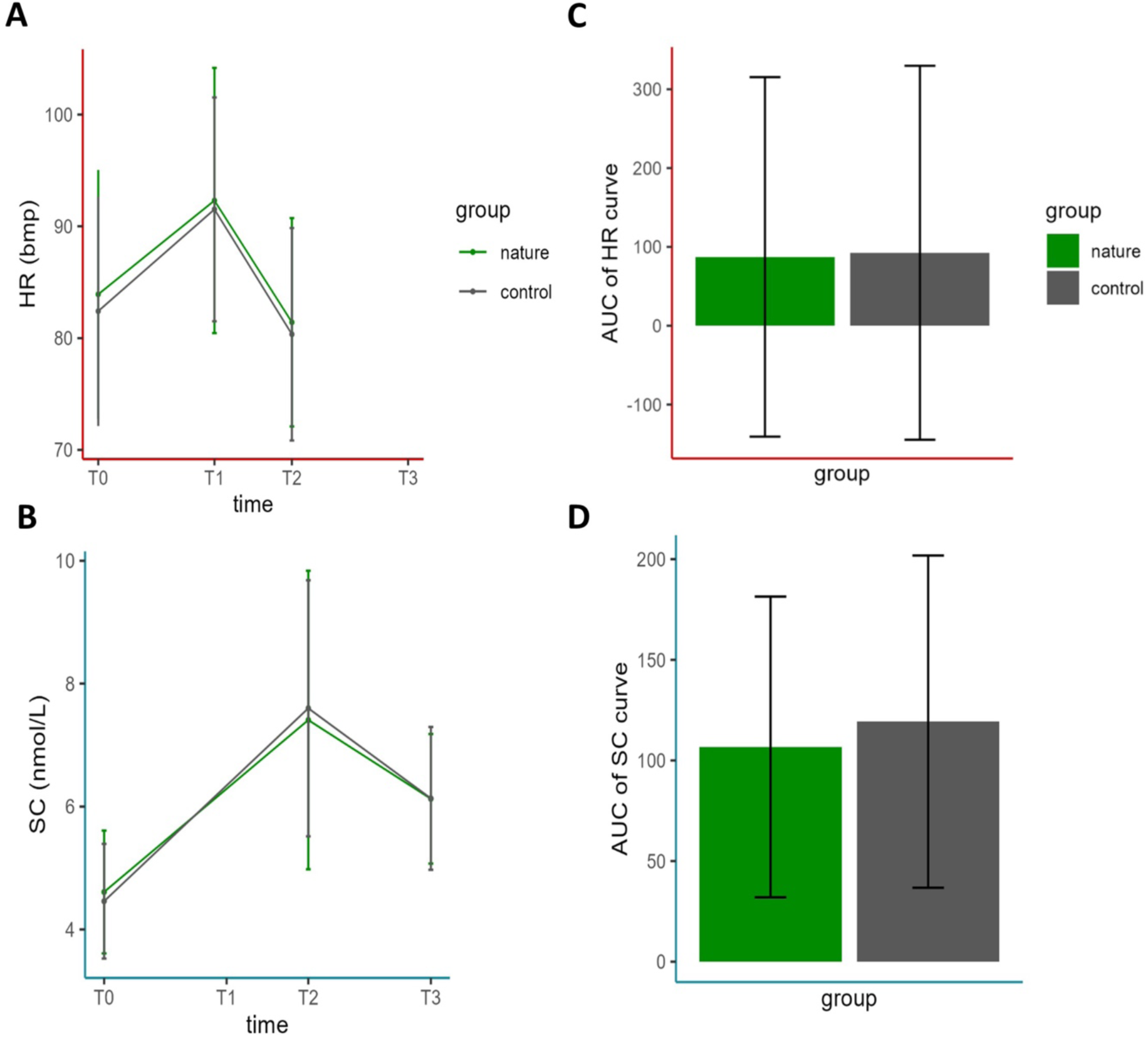
The effects of VR-simulated green space on post-stress biological responses. (A) Dynamic changes in heart rate (HR) are compared between the two groups from baseline (T0) to post-intervention (T2). (B) Between-group comparisons of the area under the curve (AUC) for HR responses. (C) Dynamic changes in skin conductance (SC) between the two groups from baseline (T0) to post-task (T3). (D) Between-group comparisons of the area under the curve (AUC) for SC responses.

Cortisol data similarly indicated a significant time effect (F(1.53, 97.97)= 68.48, *p*<.001, partial η^2^= 0.52.) without group effects (F(1,64)= 0.004, *p*=.95, partial η^2^= 0.) or interaction (F(1.53, 97.97)= 0.23, *p*=.74, partial η^2^=0.004; **Figure3B**), supported by Bayesian factors exceeding 100 (BF_10_>100). An independent samples t-test on the AUC for cortisol showed no significant group differences (t (64) =0.65, *p*=.52, Cohen’s d=0.16; **Figure3D**), indicating no VR effect on cortical responses. Bayesian t-tests further supported the hypothesis of no group differences (BF_10_= 0.30), suggesting that VR-stimulated green spaces do not mitigate biological stress responses.

### 3.3 Does VR stimulated green space prevent stress-induced working memory impairment?

We assessed changes in working memory (WM) performance between two groups to determine whether VR-stimulated green spaces could mitigate WM impairment following stress induction. Descriptive analysis results are presented in **Table2**. In the Digit Symbol Substitution Task (DSST), repeated measures ANOVA (RMANOVA) indicated a significant effect of time (F (1,64)= 35.89, *p*<.001, partial η^2^= 0.36), with no significant group differences (F (1,64)= 0.007, *p*= .93, partial η^2^= 0.), or time-group interactions (F (1,64)= 0.41, *p*= .53, partial η^2^= 0.006; **Figure4A**). Bayesian analysis supported the model that included only time (BF_10_=7.35). Similarly, in the visuo-spatial (VS) task, we observed the effect of time (F (1,60)=25.49, *p*<.001, partial η ^2^= 0.30), with no significant differences between groups (F(1,60)= 1.52, *p*= .22, partial η^2^= 0.03) or interaction (F(1,60)= 0.11, *p*= .75, partial η^2^= 0.002; **Figure4B**), supported by a BF_10_ of 4.87. The phonological loop (PL) task showed similar trends: significant effect of time (F(1,63)= 22.49, p<.001, partial η^2^= 0.26.), with no between-group effects (F (1,63)= 0.40, p= .53, partial η^2^= 0.006.) or interactions (F(1,63)= 1.10, p= .30, partial η^2^= 0.02; **Figure4C**), with Bayesian analysis yielding a BF_10_ of 6.29. Collectively, our data do not support a protective effect of VR-stimulated green spaces on WM under acute stress conditions.

**Table2.**
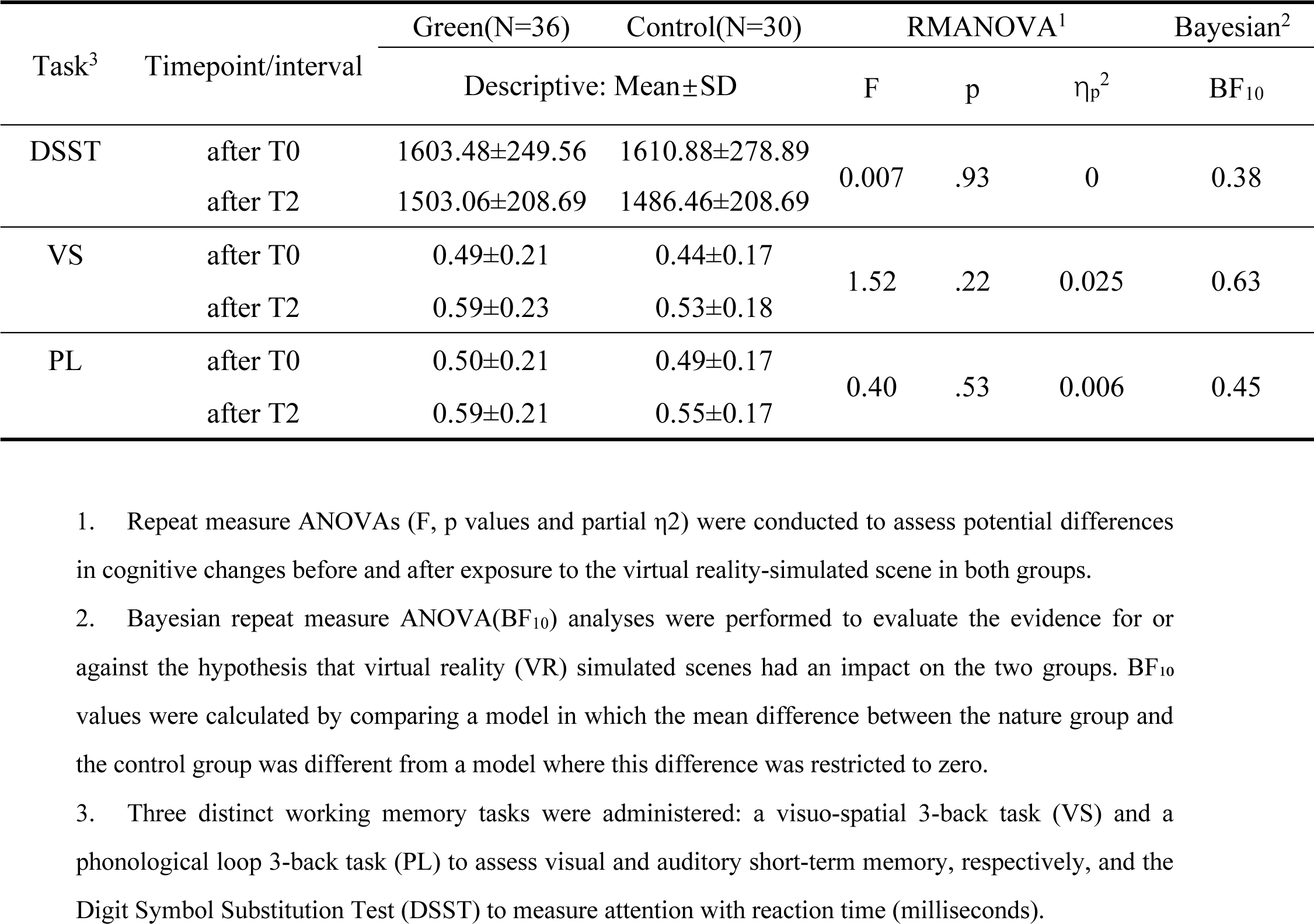
Comparison of cognitive task performance change across time between experimental group and control group.

**Figure4.**
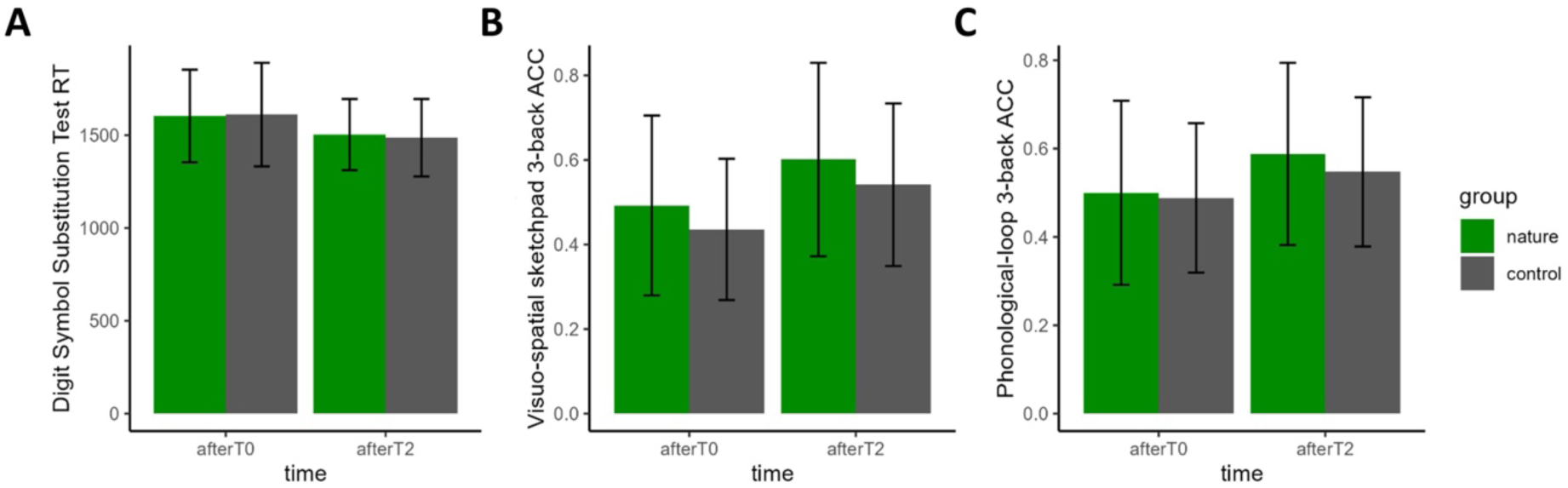
Cognitive tasks performance at the baseline (T0) and post-intervention (T2). **(A)** Performance (i.e., Reaction Time (RT) of the Digit Symbol Substitution Test (DSST). **(B)** Performance (i.e., Accuracy (ACC)) of the visuo-spatial (VS) 3-back task. **(C)** Performance (i.e., Accuracy (ACC)) of the phonological-loop (PL) 3-back task.

### 3.4 Exploratory analyses to link biological responses to stress with working memory performance

Our full-sample main analyses revealed no effect of green space virtual reality on reducing the biological response to stress or providing additional protection to working memory under stress. We conducted two exploratory analyses within subsamples to further confirm these results. First, to eliminate bias caused by the AUC estimation of nonstandard cortisol changes, we focused our exploratory analyses considering participants’ cortisol response curves. A post hoc test of cortisol levels at T0 and T3 indicated that most participants’ cortisol levels did not adequately recover (t = -6.32, *p* < .001, Cohen’s d = -1.02; M_T0_ = 4.54, M_T3_ = 6.13; **Figure 5A**). By observing individual cortisol response patterns, we identified six change modes, involving 13 participants (6 from the green group; 7 females) in the exploratory analysis. A post hoc test of this subsample indicated that cortisol levels subsided to baseline in both groups (t = -0.29, *p* = .78, Cohen’s d = -0.10; Mₜ₀ = 4.95, Mₜ₃ = 5.07). RMANOVA analysis showed no effect of VR exposure on AUC (t(11) = 0.81, p = .43, Cohen’s d = 0.45; **Figure 5B**), while Bayesian analysis provided moderate to weak evidence with a BF₁₀ of 0.57. In the DSST task, analysis found no effect of VR exposure (F(1,11) = 1.31, *p* = .28, partial η² = 0.11) or time (F(1,11) = 2.75, *p* = .13, partial η² = 0.20), nor was there any interaction (F(1,11) = 0.26, *p* = .62, partial η² = 0.02; **Figure 5C**). Meanwhile, Bayesian analysis provided weak evidence for a model where only time had an effect (BF₁₀ = 1.02). In the VS task, RMANOVA revealed a significant time effect (F(1,11) = 11.28, *p* = .006, partial η² = 0.50), but no VR exposure effect (F(1,11) = 2.50, *p* = .14, partial η² = 0.19), nor any interaction (F(1,11) = 0.27, *p* = .61, partial η² = 0.02; **Figure 5D**). However, Bayesian analysis supported considerable evidence for a model where both VR exposure and time had an effect (BF₁₀ = 8.71; M₉₀ = 0.67, M_control_ = 0.50). In the PL task, we found no time effect (F(1,11) = 2.46, p = .15, partial η² = 0.18), VR exposure effect (F(1,11) = 1.97, p = .19, partial η² = 0.15), or interaction (F(1,11) = 0.04, p = .85, partial η² = 0.003; Figure 5E). Bayesian analysis provided subtle evidence (BF₁₀ = 0.90).

**Figure 5.**
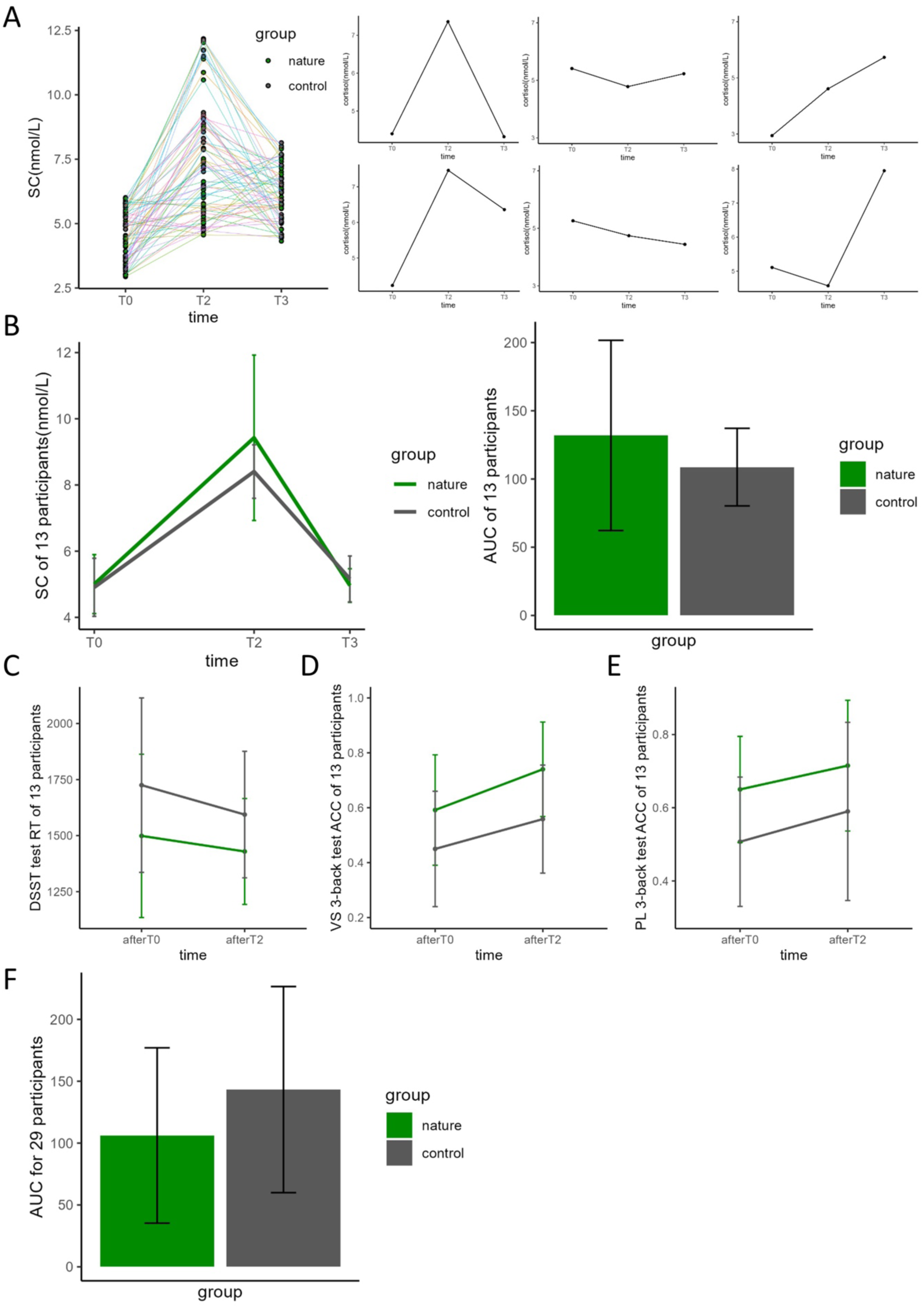
Exploratory analysis of cortisol responses and working memory performance in subsamples. (A) Salivary cortisol (SC) change patterns over time varied among individuals (left panel), categorized into six distinct types (right six subplots). (B) Cortisol responses of the 13 participants with adequate cortisol reduction. The left panel shows the SC level change curve, while the right panel displays the mean SC total response from T0 to T3 for both groups. mean SC levels at T0 and T3 for both groups. (C) Working memory performance in the Digit Symbol Substitution Test (DSST) under VR and control conditions, measured by reaction time. (D) Accuracy (ACC) in the VS 3-back task. (E) ACC in the phonological loop (PL) 3-back task. (F) AUC estimation of cortisol response curves in participants with stress-induced WM impairment in both groups.

Next, we performed a second exploratory analysis on participants who showed significant cognitive impairment under stress, selecting those with a decline in at least one task performance. Consequently, 29 participants were analyzed (16 in the green group, 26 females). Classical analysis found no significant difference between the two groups in AUC (t(27) = -1.30, *p* = .21, Cohen’s d = -0.48), while Bayesian analysis provided subtle evidence (BF₁₀ = 1.28; M_green_ = 106.15, M_control_ = 143.24; **Figure 5F**). This exploratory analysis suggests that in participants with significant stress-induced WM impairment, green space VR may help reduce cortisol responses.

## 4. Discussion

Recent advances in environmental psychology underscore the potential of green spaces to mitigate stress responses. Nevertheless, there remains no consensus on whether a brief, single-session VR simulation of green spaces can reduce acute stress responses and preserve cognitive function in humans under acute stress. Our study provides evidence against this notion, showing that the 15 minutes exposure to VR-simulated green spaces does not alleviate acute stress responses nor protect against impairments in working memory under acute stress. These findings necessitate careful methodological considerations for future stress management strategies using VR-simulated green spaces.

Restorative environment theory^51^ posits that exposure to green spaces may reduce stress, alleviate mental fatigue, and enhance cognitive function. Numerous studies report physiological benefits, such as decreased levels of plasma endothelin-1 (ET-1), cortisol, and blood pressure from exposure to natural green space. They also document cognitive and mental health improvements in routine tasks and general well-being, addressing stress, depression, and anxiety^52–61^. Remarkably, green spaces have provided effective, sustained interventions even under severe stressors like the COVID-19 pandemic, mitigating stress and anxiety among urban populations^22,62,63^. While the precise mechanisms by which nature counters the adverse effects of stressors remain partially undefined, green spaces are a promising method for managing stress in both the general population and individuals with mental disorders.

Although the mental health benefits of green space exposure are well-documented, access may be limited by various factors, particularly in urban areas devoid of such environments. Virtual reality (VR)-simulated green spaces offer a feasible, acceptable, and potentially effective short-term intervention for stress reduction^64^. Despite a strong theoretical basis for the acute stress-reduction benefits of natural environment immersion, empirical support remains provisional^65^. Contrasting with previous studies that demonstrated benefits on mental health, our study found no evidence that VR-simulated green spaces reduce acute stress responses or protect working memory performance under stress. Importantly, these studies often compared natural environments with urban settings or varied the amounts of green elements^52,56–60,66^. These distinctions might reveal the negative impacts of VR-simulated urban spaces rather than the advantages of green spaces. Our study employed a strictly matched blank control scene and uses a Bayesian analytical framework alongside traditional statistical methods to differentiate between the ‘absence of evidence’ and ‘evidence of absence’ of VR effects. Moreover, this study is the first to design an interactive green VR scene where participants could ‘walk’ and ‘touch’ objects, unlike previous settings where participants were stationary and passively observed the environment—analogous to real-world recreational control. We hypothesized that this interactivity would enhance the mental benefits of green spaces within a VR context. However, our comprehensive analysis found no evidence to support the benefits of VR-simulated green spaces on stress responses or working memory performance when compared to the control scene.

The absence of significant benefits from VR-simulated green space on biological stress responses and working memory performance may be due to the specific design of our VR scenes. To achieve more precise experimental control, our scenes lacked interactive elements, such as touching plants or climbing hills, and omitted components like terpenes and air pollutants ^17^. The static nature of the VR scene also introduced potential stressors, including an eerie silence and solitude reminiscent of the ‘uncanny valley’ effect ^67^, eliciting feelings of dread and loneliness among some participants. Second, our VR protocol consisted of a single 15-minute session, whereas prior research emphasizes the importance of the living environment and daily exposure for mental health benefits^22,53^. We recommend that researchers openly share VR scenes that have proven beneficial in small-scale studies to facilitate direct replication, assess the effects of active ingredients, and support long-term, multi-session interventions.

### Limitations and future directions

We acknowledge several limitations of our study and propose directions for future research. The first limitation concerns participant demographics. Although the gender ratio was statistically balanced between the experimental and control groups, approximately 84% of our participants were female. Previous laboratory studies on acute stress predominantly recruited male participants to avoid confounds related to gender differences and menstrual cycle-dependent variance in stress responsiveness ^68^. Variability in cortisol levels may necessitate a larger sample size to detect significant experimental effects. Future studies could use a gender-balanced sample to detect potential gender-specific effects and potential boundary conditions that certain group of participants could benefit from the VR green space intervention. Additionally, our sample comprised mentally healthy university students, which limits the generalizability of our findings to other populations with poorer mental health or higher baseline stress levels (e.g., individuals with stress-related psychiatric disorders such as depression, anxiety, and PTSD), despite the potential efficacy of the VR intervention for these groups.

A second limitation is that both the experimental and control groups received VR interventions—one group was exposed to a green space, and the other to a gray, empty scene— without a proper additional control group that did not receive any VR intervention. This omission hinders our ability to accurately estimate the effect size of stress-induced working memory (WM) impairment. Although the effect of acute stress on WM is well-established in previous studies^8–10,16^, it remains crucial to disentangle the effects of stress and VR intervention in our study. It is possible that the gray empty condition may have reduced stress responses and protected WM for some participants, suggesting that both VR conditions may have mitigated stress-induced WM impairment. Future research could experimentally test this hypothesis.

The third limitation concerns the ecological validity of our VR program. Although we made every effort to replicate the visual features (primarily the visual presentation of greenery and plant shapes) and activities, such as walking and interacting with plants, other essential aspects of walking in a natural green space remain unaddressed. Key elements, such as odors, sounds, the intensity and pattern of aerobic exercise, and social-affective factors (e.g., walking with friends and family), all play crucial roles in generating the beneficial effects associated with natural green environments^17^. Our study suggests that visual presentation alone may not be sufficient to achieve these effects. Future research should systematically investigate the contribution of various elements of walking in green spaces and their interactions. As VR technology continues to develop, it may eventually be possible to incorporate all these factors simultaneously into VR programs, thereby achieving optimal intervention effects.

## 5. Conclusions

A growing body of research supports the mental health benefits of exposure to green spaces, particularly in reducing stress responses. In our study, we used VR technology along with various working memory paradigms to assess whether simulated green space exposure following acute stress induction could attenuate biological stress responses and prevent stress-induced impairment in working memory. We created two meticulously matched VR scenes, including a gray empty scene as a control. By continuously monitoring heart rate and cortisol levels post-stress induction, we validated the effectiveness of our stress induction protocol. Contrary to our initial hypotheses, the findings indicated that, compared to the control VR setting, the green space environment did not provide additional benefits in reducing biological stress responses or in shielding working memory from stress. Collectively, our results question the widely held view that VR green spaces can effectively substitute for actual green spaces in managing stress. Given the escalating interest in leveraging green space exposure for mental health benefits, our findings advise caution against further investments before the impacts of VR-simulated natural and urban spaces on human mental health are fully understood.

## Acknowledgments and Funding

This study was supported by the National Natural Science Foundation of China (grant No. 32300879), Humanities and Social Sciences Fund, Ministry of Education (grant No. 22YJCZH109), Natural Science Foundation of Hubei (grant No. 2022CFB793), Open Research Fund of the State Key Laboratory of Cognitive Neuroscience and Learning (CNLYB2103), Open Research Fund of the Key Laboratory of Adolescent Cyber Psychology and Behavior (grant No. CCNUCYPSYLAB2022B10), the Major Program of the National Social Science Foundation of China (grant No. 22&ZD187), self-determined research funds of CCNU from the colleges’ basic research and operation of MOE(CCNU22QN020), and National Natural Science Foundation of China (62077010)

## Contributions

W.L. H.Z and Y.Y.X conceived the Study; Q.Y.Y, X.X.Z, Z.X.Y, Y.Y.X and G.H.D collected the data; W.L. and J.S.L analyzed the data; W.L., H.Z and Y.Y.X prepared the first draft. W.L., H.Z and Y.Y.X reviewed and edited the manuscript. W.L and H.Z provided supervision and obtained funding.

## Supplementary Materials

**Supplementary Table 1.**
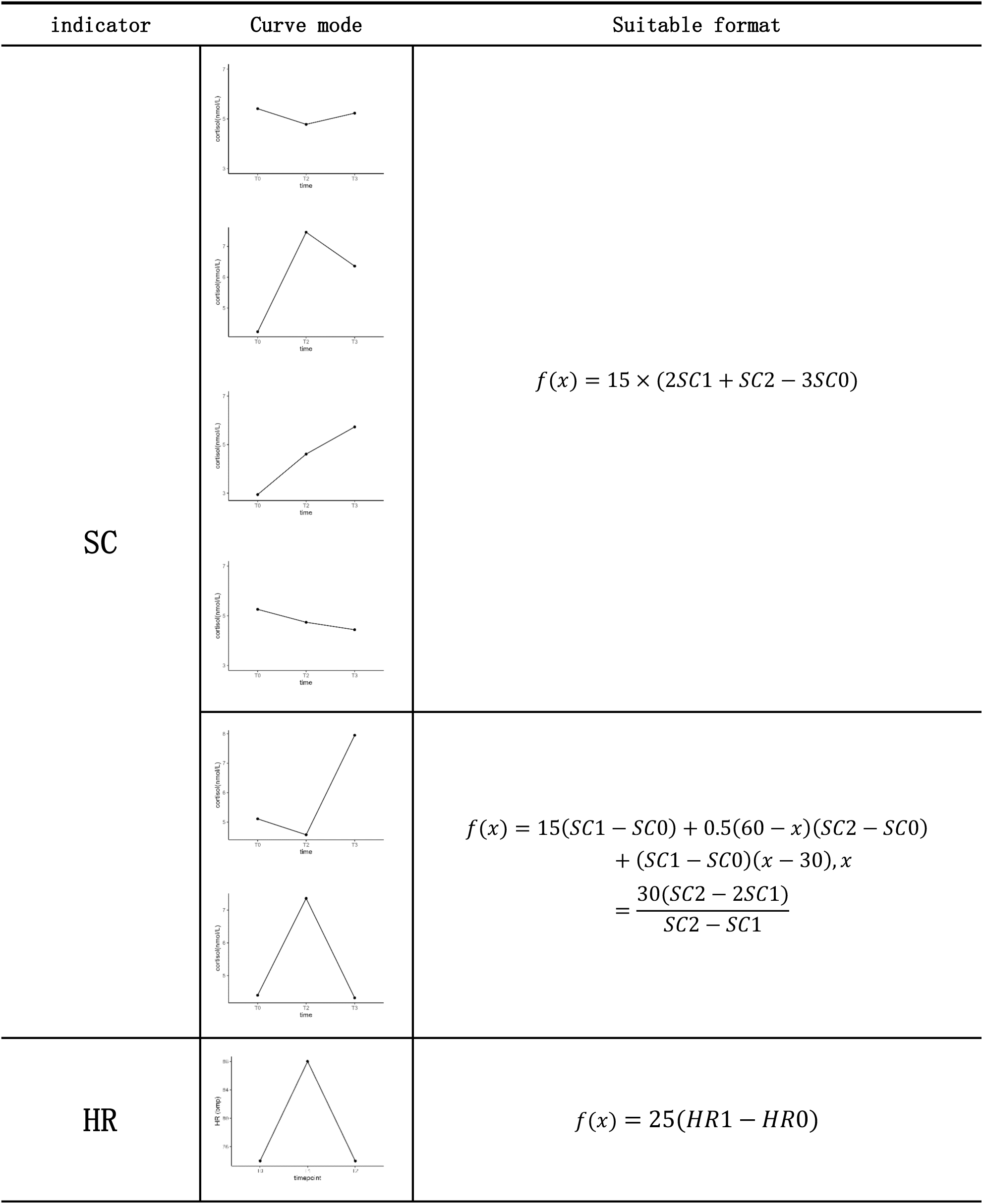
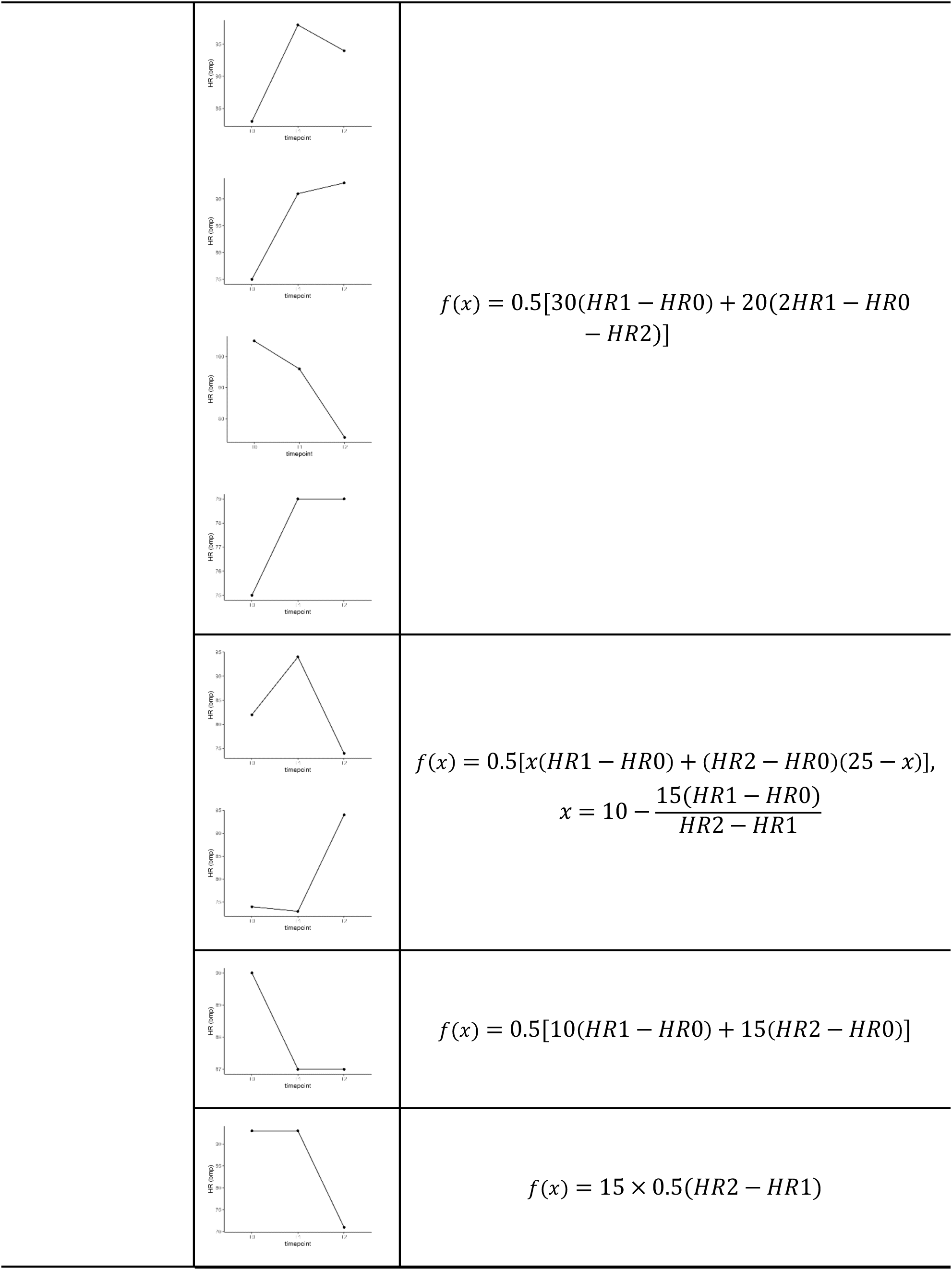
The cortisol change mode and matched format.

### AUC calculation depended on its curve mode

Since the AUC is a vector rather a scalar, we deemed y=SC0 as standard line, crossing point (SC2, T3) to it as a perpendicular line and dividing the closed figure into two parts. The AUC was defined as the sum of the two vector parts’ area, whose value could be negative. See Supplementary Table 1.

### Participants selection rules

In the explore analysis, we first selected 13participants from cases in Figure5 A left-upper and those difference between SC2 and SC0 was smaller than 1 nmol/L (namely measurement error) in Figure5 A left-down of 6 pictures. We checked VR-stimulated scene effect based on these 13 participants data.

We operational defined cognitive damage as at least a sort of tasks performance declined, consequently reaping 29 cases. It seems that there’ s no good reason to compare the difference of repeated cognitive tasks performance, since we were aiming at the difference of AUC and the definition of cognitive damage leaving the participants performed poorly in one task might do well in other tasks. But at here, we’ d like to show the explore analysis result using 29 participants’ cognitive tasks data. Repeat measure ANOVA (RMANOVA) showed main effects of time (F(1, 27)= 5.51, p= .03, partial η^2^= 0.17.) and group (F(1,27)= 2.54, p= .12, partial η^2^= 0.09.) and no interaction (F(1, 27)= 0.56, p= .46, partial η^2^= 0.02.) in DSST task. Bayesian analysis faintly supported the model where only time had an effect with BF_10_=1.97. RMANOVA found no significant main effect in VS task (group: F(1, 23)= 0.03, p= .88, partial η^2^= 0.00; time: F(1,23)= 1.63, p= .22, partial η^2^= 0.07.), neither nor an interaction (F(1,20)= 0.01, p=.92, partial η^2^= 0.00.). Bayesian analysis supported no effect model. In PL task, the same result recurred (group: F(1, 27)= 0.09, p=.76, partial η^2^= 0.00; time: F(1, 27)= 0.64, p=.43, partial η^2^= 0.02.), but a marginal significance in interaction (F(1, 27)= 3.45, p=.07, partial η^2^= 0.11.). However, Bayesian analysis supported no effect model. The mean of two group showed an apt divergence in two groups, namely increased performance in nature group with a reduction in control group. See Supplementary Table 1.

**Supplementary Table 2.**
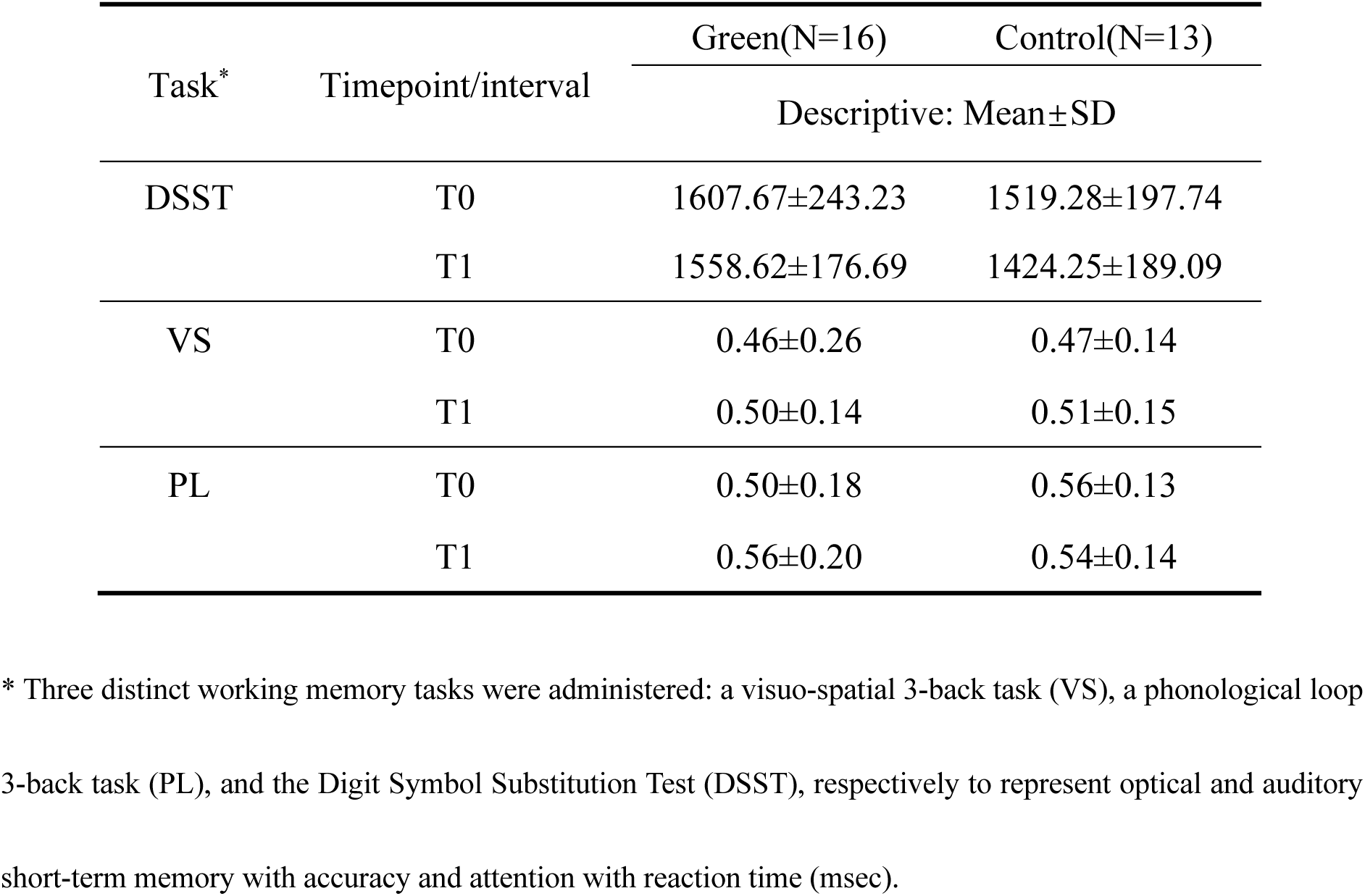
Descriptive analysis result of cognitive task performance repeated measurement in two group.

| Task* | Timepoint/interval | Green(N=16) | Control(N=13) |
| --- | --- | --- | --- |
|  |  | Descriptive: Mean±SD |  |
| DSST | T0 | 1607.67±243.23 | 1519.28±197.74 |
|  | T1 | 1558.62±176.69 | 1424.25±189.09 |
| VS | T0 | 0.46±0.26 | 0.47±0.14 |
|  | T1 | 0.50±0.14 | 0.51±0.15 |
| PL | T0 | 0.50±0.18 | 0.56±0.13 |
|  | T1 | 0.56±0.20 | 0.54±0.14 |

\* Three distinct working memory tasks were administered: a visuo-spatial 3-back task (VS), a phonological loop 3-back task (PL), and the Digit Symbol Substitution Test (DSST), respectively to represent optical and auditory short-term memory with accuracy and attention with reaction time (msec).

